# Terrestrial laser scan metrics predict surface vegetation biomass and consumption in a frequently burned southeastern U.S. ecosystem

**DOI:** 10.1101/2023.01.15.524107

**Authors:** E. Louise Loudermilk, Scott Pokswinski, Christie M. Hawley, Aaron Maxwell, Michael Gallagher, Nicholas Skowronski, Andrew T. Hudak, Chad Hoffman, J. Kevin Hiers

**Affiliations:** USDA Forest Service, Southern Research Station, Athens Prescribed Fire Laboratory, Athens, GA, USA; New Mexico Consortium, Center for Applied Fire and Ecosystems Science, Los Alamos, NM, USA; West Virginia University, Department of Geology and Geography, Morgantown, WV, USA; USDA Forest Service, Northern Research Station, Climate, Fire, and Carbon Cycle Sciences, Morgantown, VA, USA; USDA Forest Service, Northern Research Station, Climate, Fire, and Carbon Cycle Sciences, Silas Little Experimental Forest, New Lisbon, USA; USDA Forest Service, Rocky Mountain Research Station, Forestry Sciences Laboratory, Moscow, ID, USA; Colorado State University, Warner College of Natural Resources, Fort Collins, CO, USA; Texas A&M University, Natural Resources Institute, 1747 Pennsylvania Ave, NW, Suite 400, Washington, DC, USA

## Abstract

Fire-prone landscapes found throughout the world are increasingly managed with prescribed fire for a variety of objectives. These frequent low-intensity fires directly impact lower forest strata, and thus estimating surface fuels or understory vegetation is essential for planning, evaluating, and monitoring management strategies and studying fire behavior and effects. Traditional fuel estimation methods have applications for stand-level and canopy fuel loading; however, local-scale understory biomass remains challenging because of complex within-stand heterogeneity and fast recovery post-fire. Previous studies have demonstrated how single location terrestrial laser scanning (TLS) can be used to estimate plot-level vegetation characteristics and impacts from prescribed fire. To build upon this methodology, co-located single TLS scans and physical biomass measurements were used to generate linear models for predicting understory vegetation and fuel biomass as well as consumption by fire in a southeastern U.S. pineland. A variable selection method was used to select the six most important TLS-derived structural metrics for each linear model, where model fit ranged in R^2^ from 0.61 to 0.74. This study highlights a prospect for efficiently estimating vegetation and fuel characteristics relevant to prescribed burning via the integration of a single-scan TLS method adaptable by managers and relevant for coupled fire-atmosphere models.

## Introduction

Frequently burned ecosystems are found throughout the world and are known for their role in supporting high levels of biodiversity and structurally complex understory plant communities [1-3]. The architectural structure of these species-rich plant communities is characterized by fine-scale (<1 m) heterogeneity that has evolved with low-intensity surface fires. Prescribed burning is used to maintain these fire-dependent ecosystems, and practitioners monitor and estimate surface fuels to gauge the appropriate fire frequency and formulate prescribed fire management strategies [4,5]. Furthermore, characterization of the surface fuels are a critical input for a range of fire behavior and effects models (e.g. BehavePlus, QUIC-Fire, FIRETEC and the Wildland Urban Interface Fire Dynamics Simulator)[6-10]. At the stand scale (10’s to 1000’s of ha), approximate fuel loading or biomass estimates are dependent on ecosystem type, burn history, and plant species composition as well as climate and soil characteristics that influence productivity and ultimately fuel accumulation rates [1,11]. Current datasets on surface vegetation or fuel biomass are available, but only provide estimates in horizontal dimensions, without a vertical dimension, and at spatial resolutions ranging from 30-m resolution satellite imagery to stand-level averages for representative ecosystems [11,12]. Though these approaches have been successfully used by managers in a variety of situations including large-scale fire management, they do not provide information on the fine scale variability in fuels that drive fire behavior and effects in prescribed fires and in frequently burned ecosystems. There are no operational products that provide information on finer-scale attributes of surface vegetation composition and architecture, despite research in laboratory, field, and modeling environments that have identified these biophysical characteristics as drivers of variation in fire behavior and effects from surface fires [13-17]. Thus, improving estimation of the spatial resolution of fuels and including information about the vertical dimensionality of the fuelbed architecture or 3D characteristics of surface live and dead vegetation are critical to improving fire behavior predictions for surface fires.

The spatial variation of biomass at scales relevant to surface fire is the most difficult to capture because of the complexity in composition and distribution within and across stands, even within the same ecosystem and same time since last burn event [6,15,18]. In southeastern U.S. ecosystems where production and turnover rates are high [19,20], repeated burning provides a continuum of fine surface fuels and live vegetation that enable a broad range of fire behavior throughout the year [21,22]. As such, the low (often <1m height) vegetation (shrubs, grasses, forbs, leaf litter, soil organic layer, and small coarse wood) continuously changes with each fire and ecological response [13], making estimates of surface vegetation structure and loading, notwithstanding fuel moisture dynamics [23], highly variable at fine temporal (<hr) and spatial scales (<1m). Furthermore, large diameter coarse woody debris (>20 cm) have little opportunity to accumulate and hence contribute little to combustion during prescribed burns because of their high moisture retention and fast decay rates [24,25]. Furthermore, consumption, or the amount of combusted biomass defines a fire’s intensity, and as such is used as a proxy for estimating fire effects [26]. The complex fuelbed and fine-scale fire-atmosphere interactions during a prescribed fire cause highly variable consumption patterns across a burned area. Given these ecosystem processes and combustion patterns, monitoring of these areas is needed as often as burning occurs, yet current approaches fall short of estimating heterogeneous biomass change and consumption within and across stands.

While remote sensing technology, particularly terrestrial laser scanning (TLS), can capture this fine-scale 3D structure, surface biomass variation is more difficult to characterize because mass varies more by fuel type than by its shape or volume [27-29]. To address this, approaches have been developed to couple traditional plot level measurements of mass with TLS data. Over a decade ago, Loudermilk et al. (2009) illustrated that traditional estimates of volume (sphere, cylinder) significantly overestimates the volume of vegetation, and that fine-scale volume estimated from TLS (voxel-based method) can be linked directly to fine-scale leaf area and mass of two common southeastern shrub species. Since then, field sampling has transitioned from 2D to 3D (Hawley et al. 2018) and examples of using TLS to characterize understory vegetation while linking to estimates of biomass or leaf area have increased [28,30-33].

Many of these earlier studies using TLS used research grade instrumentation and processing techniques that require specialized software and knowledge, which are time consuming and not suitable for management needs. Fortunately, recent advances of lidar technology has allowed for a transition from research to management applications, specifically through available low-cost, portable push button instruments [34,35]. Instrument types range from hand-held or vehicle mounted scanners [36,37], unmanned aerial vehicle scanners [38], stationary scanners [39-41], and even mobile phone apps [42]. These all vary in quality, accuracy, and inherent laser capabilities (range, output point density, number of returns) that all impact each desired forest measurement attribute [34,43]. While these instruments are now affordable (<$30,000), widely available, and relatively easy to use, manipulating, merging multiple scans, and analyzing the output 3D point clouds still requires specialized software and high-level coding and analytical skills.

There is growing interest to streamline this analytical bottleneck by capitalizing on single location terrestrial laser scans to characterize plot-level fuel and vegetation characteristics for use in ecosystem and fire effects monitoring and to provide for spatially explicit fire behavior and effects models. A new method, developed by Pokswinski et al. (2021) was spearheaded in the southeastern U.S., where prescribed fire is the prevailing forest management tool. This TLS-based approach of characterizing understory vegetation capitalizes on the data richness of a single scan, with typically ∼6-8 million 3D points, measures nearly the entire 3D structure of a small, forested area within five minutes [44]. For comparison, these plots, clipped to a 15m radius, are equivalent to a typical forestry plot size of 0.07 ha (0.18 ac), where a portion of that area is measured by hand (e.g.,[45,46]). Currently, over 100 metrics, or structural characteristics can be calculated from each point cloud, which represent the vegetation’s spatial distribution, density, proportion or identification of various parts of the forest (e.g. tree boles) as well as the structure of openings or space within the forest, and differences between true empty space and space created by occlusion. These metrics have been successful in predicting fire severity [47], forest structure [48], and understory species richness [40], but have yet to predict surface vegetation mass or consumption. The difficulty lies in the laboratory processing, which requires time and resources to sort, dry, and weigh shrubs, grasses, leaf litter and woody debris collected *in situ*, and analytical complexity is relating 3D structure to mass. As such, there is a need to develop an approach to easily predict the heterogeneity in surface vegetation biomass across ecosystems with heterogeneous vegetation structure and composition.

The objective of this study was to build upon the most recent methodology for developing vegetation metrics from single location terrestrial laser scans [44] by linking surface biomass to these metrics. We assessed the relationship of eight vegetation and fuel classes of surface biomass to 162 TLS vegetation and fuel metrics in a frequently burned longleaf pine (*Pinus palustris*) woodland in southeastern Georgia, USA. We used a robust variable selection method to develop linear models for each of these eight classes. We used these linear models to estimate pre-burn mass, post-burn mass, as well as mass consumption by fire.

## Methods

### Study site

Fort Stewart - Hunter Army Airfield (113,017 ha, 31°56’ N, 81°36’ W, elevation 2-56 m) is primarily located in Liberty and Bryan counties in the Coastal Plain Province on the southeastern Atlantic coast of Georgia [49]. This region has hot and humid summers with short, mild winters with average monthly temperatures ranging from 27.92 C in July to 11.06 C in January (NCEI Climatology 1991-2020 dataset, Fort Stewart/Wright, GA US Weather Station, 31.869 deg N, 81.624 deg W). The average annual rainfall is 1298 mm (NCEI Climatology 1991-2020 dataset, Fort Stewart/Wright, GA US Weather Station, 31.869 deg N, 81.624 deg W). Fort Stewart falls within the historical range of the longleaf pine ecosystem (Frost 2006), which was severely reduced by logging in the 19^th^ and early 20^th^ centuries. Prior to the federal acquisition of this land in the 1940s, the landscape was fragmented by agricultural fields [50,51]. Currently, the overstory of these continuous sandhills and flatwoods is dominated with restored longleaf pine (*Pinus palustris*), slash pine (*P. elliottii*), loblolly pine (*P. taeda*), and turkey oak (*Quercus laevis*). The understory is a diverse mixture of graminoids, forbs, and shrubs such as wiregrass (*Aristada stricta*), gallberry (*Ilex glabra*), and saw palmetto (*Serenoa repens*). Beyond military missions, the land is managed for a variety of plant and animal species, including the endangered Red-cockaded Woodpecker (*Picoides borealis*), regular timber harvesting and year-around prescribed burning on a 3–5 year interval is used to maintain desired canopy composition and density, as well as reduce fuels loads [52-54]. Our study site at Fort Stewart consisted of two adjacent burn units – F6.3 (261 ha) and F6.4 (397 ha) that was represented by longleaf pine flatwoods and intermixed cypress wetlands (*Taxodium* spp.). These burn units are on Pelham loamy sand, Leefield loamy sand, and Ellabelle loamy sand which are very deep, somewhat poorly to very poorly drained, moderately slow to moderately permeable soils that formed in unconsolidated Coastal Plain sediments [55]. These areas had not been burned in three years; the last prescribed burn occurred in 2019.

These two burn units were burned on two consecutive days, March 2nd and 3rd, 2022. During February 2022, the area received 29.48 mm of precipitation with the last event of 5.84 mm occurring on February 28 (Fort Stewart/Wright, GA US Weather Station, 31.869° N, 81.624° W). On the burn days, average direction was west to northwest (284.30493 and 260.7581) and average wind speed was 4.52 kph with maximum speeds of 12.96 kph and 22.22 kph respectively (Fort Stewart/Wright, GA US Weather Station, 31.869 deg N, 81.624 deg W). The mean temperature was 16.06 C° and relative humidity ranged from 16% to 89%; averaging 53% (Fort Stewart/Wright, GA US Weather Station, 31.869 deg N, 81.624 deg W).

### Sampling Design, Data Collection, and Processing

In February 2022, forty-one plots were randomly placed throughout the longleaf pine Flatwoods of the two chosen burn units at Ft. Stewart (Figure 1). The interior wetlands were not sampled as they are often inundated with water and as such typically do not ignite or contribute to fire spread. First, one lidar scan was collected in the center of each plot using a Leica BLK360 (Leica Geosystems, Heerbrugg, Switzerland). More details on the laser system and processing are in the “Terrestrial Lidar Processing” section below. Approximately three meters from the plot center and at a 45° azimuth, a pre-burn clip plot was established. A post-burn clip plot was also established within three meters from the plot center. This plot was visually identified to have similar fuel composition and structure to the pre-burn clip plot. This provided for a paired sampling approach to estimate consumption, where pre-burn mass (clipped and removed) can be linked to residual mass of similar vegetation and fuel types [56,57].

**Figure 1.**
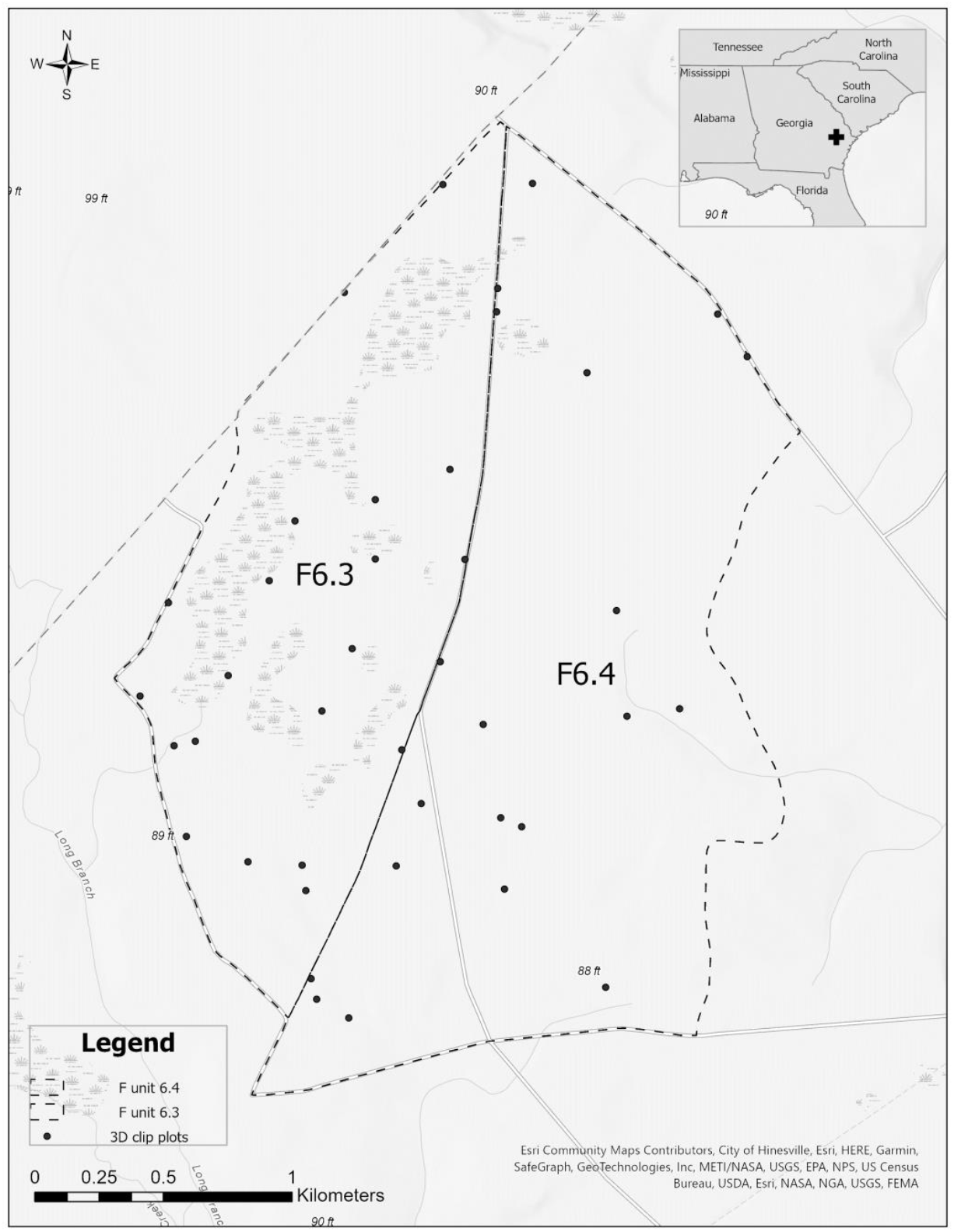
Study area and sampling distribution of co-located TLS scans and vegetation and fuel biomass samples (n=41). They were sampled before and after two prescribed burns.

Before each burn, vegetation and fuel categories were recorded in both the pre- and post-burn clip plots. Vegetation and fuel category and biomass data were collected using a simplified approach from Hawley et al. (2018). The sampling area was 0.5m in width by 0.5m in length by 1m in height. The frame was subdivided into two vertical sampling layers or strata: ground to 30 cm and 30 to 100 cm. The vegetation and fuel categories for this site were defined as woody live vegetation, now dead woody vegetation, woody litter, woody dead and downed 1-hour fuels, 10-hour fuels, 100-hour fuels, 1000-hr fuels, pinecones, conifer litter, conifer needles, and herbaceous vegetation, which includes graminoids, forbs, and vines. The ‘now dead woody vegetation’ category was used only in post-burn sampling to classify pre-burn woody live stems that were partially consumed by the prescribed fire and the aboveground plant was clearly dead or top-killed. There were no 1000-hour fuels found in our plots. Detailed description of each vegetation and fuel category is in Appendix A. Before the burn within both the pre- and post-burn plots, presence and absence of each vegetation and fuel category was recorded within each stratum. At the pre-burn plot, biomass was destructively harvested from each stratum. After the burn within each post-burn clip plot, presence and absence of each vegetation and fuel category was recorded again and biomass destructively harvested within each stratum. In the Athens Prescribed Fire Laboratory, Athens, GA, USA, biomass was sorted, dried, and weighed to determine the dry weight (in g m^-2^) of each vegetation and fuel category. The sorted biomass was dried at 70°C until the weight of the sample no longer changed, typically within 48-72 hours. Dry mass values are found in Appendix B.

### Vegetation and Fuel Mass Classes

For this study, we used our aforementioned field-derived vegetation and fuel categories (Appendix A) to create our vegetation and fuel mass classes used for our linear modeling (Figure 2). This included total surface biomass (‘Total’), Total without fine woody debris (‘Total no FWD’), Total below 30 cm height (‘Total 0-30cm”), fine vegetation and fuels only (‘Fine Fuels’), fine woody debris only (‘FWD’), and Total below 30cm height without FWD (‘Total 0-30cm no FWD’). Total was the mass of all live and dead vegetative material. Total no FWD was the Total category minus any fine woody debris in the 10-hr and 100-hr fuel categories and pinecones. Total 0-30cm was Total mass found under 30cm in height. Fine Fuels included all grass, forb, and vine material (live and dead), dead leaves (pine and cypress needles, and broadleaves), other fine tree litter (bark flakes, reproductive organs: catkins, etc), and 1-hr fuels. Total no FWD and Fine Fuels categories were created because coarse (1000 hr) and fine woody debris (10, 100hr) and live stems are rarely consumed, or only partially consumed in these systems (Ulyshen et al. 2018), and fire practitioners often focus on “fine fuels” that are most available to burn during prescribed fire operations in the southeastern U.S. [58,59]. In fact, FWD was only 14% of the total mass, and there was no difference in FWD mass between the pre-burn (mean: 130.0 g m^-2^, std: 154.8 g m^-2^) and post-burn (mean: 127.6 g m^-2^, std:170.4 g m^-2^) samples (t-test, *p*=0.92). The Total 0-30cm category was created because most of the mass (95%) was found in this layer, and we wanted to test its predictive power compared to the entire area measured up to 100cm. In the end eight vegetation and fuel mass classes were created for our linear modeling (Figure 2). All pre-burn data was used for the first six classes (‘Total’, ‘Total no FWD’, ‘Total 0-30cm’, ‘Fine Fuels’, ‘FWD’, ‘Total 0-30cm no FWD’). The next two included the post-burn total surface biomass (‘Total 0-30cm Post’) as well as pre- and post-burn total surface biomass below 30cm combined (‘Total 0-30cm Pre & Post’).

**Figure 2.**
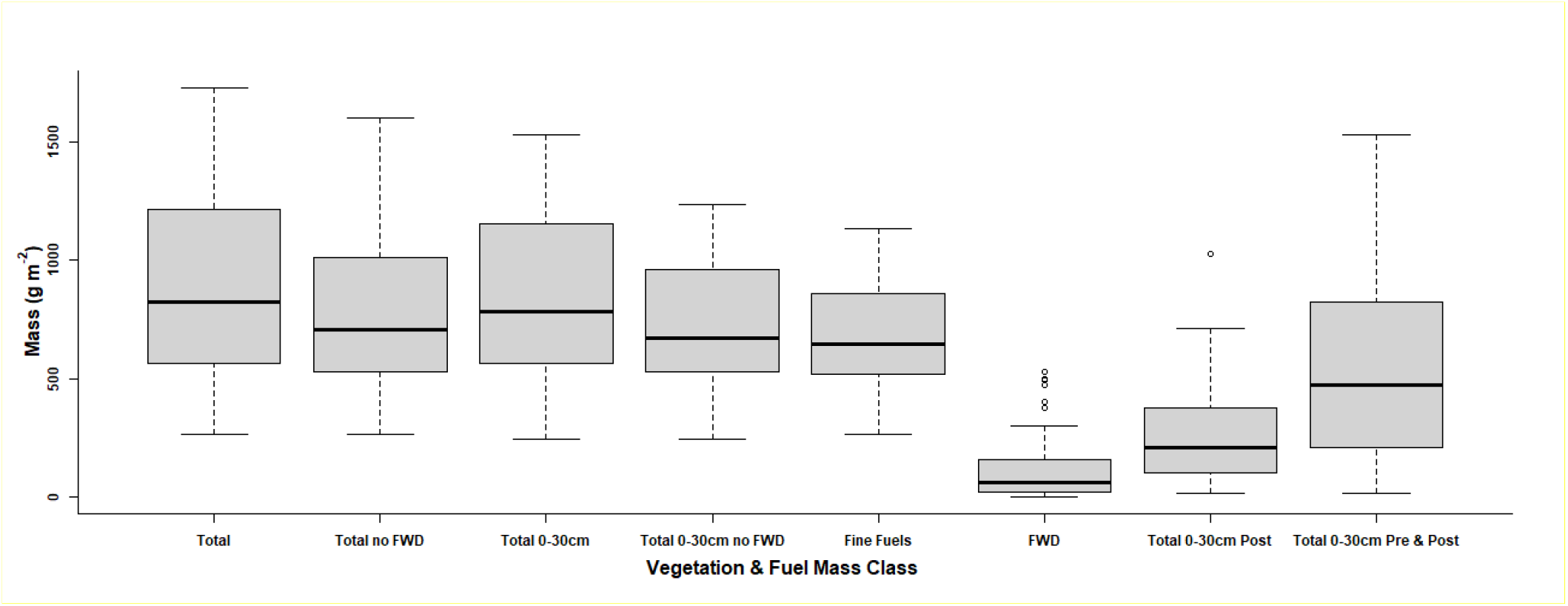
Box plots of surface dry mass (g m-2) distinguished by our vegetation and fuel mass classes. All box plots include only pre-burn biomass data (n=41), except for Total ‘0-30cm Post,’ which includes only post-burn data (n=41) and ‘Total 0-30cm Pre & Post,’ which includes both pre and post burn data (n=82). See text and Appendix A for fuel class and category descriptions.

### Terrestrial Lidar Processing

For this study, we used the Leica BLK360 (Leica Geosystems, Heerbrugg, Switzerland) terrestrial laser scanner, deployed as described above and in Pokswinski et al (2021). The BLK360 is particularly suited for forest mensuration applications because it is lightweight (1kg), affordable (<$20K), quick (< 5 minutes per scan), is splash resistant and is a single return laser. The area of coverage is 360° horizontally and 300° vertically. The laser has a wavelength of 830 nm, beam divergence of 0.4m rad, range accuracy of 4 mm at 10 m distance and 7 mm at 20 m distance, a maximum pulse rate of 360,000 points s^-1^, and maximum range distance of 60 m [39]. The unit has multiple sampling density settings to control the number of points, which can help manage data collection time and size. We used the medium density setting, which resulted in a scan time of less than four minutes.

From each single scan, a variety of metrics were calculated to summarize the point cloud and serve as predictor variables for potential inclusion in the models. Prior to this, a series of pre-processing steps were applied. The scans were first exported from the sensor to PTX format using the Cyclone Register 360 (BLK Version) software (Leica Geosystems, Heerbrugg, Switzerland). This format retains all gap values, i.e., pulses that do not have an associated return [39]. The PTX point cloud was also converted to LAS format and clipped to a 15 m radius. Then a distance-based noise filter was used to remove stray points in the point cloud. Next, the ground points were classified using a cloth simulation filter, where all ground points were normalized relative to the ground, and ground points were removed. The cloth simulation filter ground classification method classifies ground returns by modeling a rigidness-constrained cloth surface defined by an inverted point cloud [60].

A set of metrics were calculated from the PTX data to characterize the proportion of transmitted pulses that are returned. Within defined height bins (0.5, 1, 1.5, 2, >2m) and using only the non-ground points in the normalized point cloud, summary metrics for the height measures were generated to characterize the central tendency and variability of the height measurements including maximum and minimum height, mean and median height, skewness, and kurtosis. We also implement a summarization method aimed at characterizing the areas of occlusion (i.e., all pulses are returned before reaching the current volume), which makes use of spherical voxels and allows for estimating the proportion of pulses passing through a volume that is returned from that volume. Once this method is applied, summary metrics are generated from the results based on height strata, similar to the summarization process for the normalized data.

To generate additional metrics, the point cloud is also voxelized at an 8 cm^3^ spatial resolution. Trees are then segmented and stems are classified using the TreeLS package [61], and basal area, mean DBH, mean tree height, max tree height and canopy base height are calculated for all detected trees above 4cm DBH. The points classified as “stem” are then removed from the point cloud and a surface vegetation and fuels point cloud is generated with the remaining points between 0 and 3 m in height. The surface vegetation point cloud is then filtered with nearest neighbor values using the ‘fastpointmetrics’ function in TreeLS. These points are filtered using threshold values of linearity, verticality, planarity and 2-dimensional eigenratio. The result is point clouds that contain points that represent a high probability of fine fuels or coarse woody debris.

In summary, this process resulted in 162 TLS metrics, broken down by 1) how the scan was divided or not [entire scan, by stratum (0.5, 1, 1.5, 2, >2m), surface vegetation (0-3m), classified fine and coarse woody debris (0-3m)], 2) metric type (general, height statistic, quantiles, occlusion, trees), and 3) whether the metric uses the point cloud or voxelized point cloud. A detailed description of each metric is found in Appendix C.

### Linear Modeling

We applied the leaps and bounds linear regression approach, using the leaps package [62] in the R Statistical Software (v4.0.2 [63]). This approach was originally developed by [64], which was applied in the field of Forestry to test the linear relationships of a large multivariate dataset using a parsimonious approach. It performs a comprehensive search for the best subsets of variables using a ‘branch-and-bound’ algorithm and stopping rule to decide how many variables to use. If this linear regression approach proves useful, its application across similarly structured ecosystems would be less computational expensive than more complex approaches (non-linear, polynomial, etc). Linear models in R are developed using QR decomposition to solve for least squares.

Eight linear models were run using the vegetation and fuel mass classes as dependent variables and the 162 TLS metrics as independent variables. We used the ‘exhaustive’ method within the ‘subregs’ function, which performs an exhaustive approach on all combinations of variables given a target maximum subset size, in this case six. A target of six was set because we wanted to optimize the number of dependent variables for model interpretability and computational efficiency, while still cycling through all possible variable combinations. The six chosen variables were input variables to the ‘lm’ function in R to provide linear model output statistics, specifically R^2^, *p*-value, and RMSE for each model. Schwarz information criterion (BIC) [65] values were obtained from the ‘which.min’ function set for BIC generated in the ‘subregs’ function in the leaps package. Models were optimized to have low BIC values and to have the fewest variables.

Consumption was estimated using the Total 0-30cm data. Observed consumption was calculated by subtracting the post-burn mass values < 30cm in height from the pre-burn mass values <30m in height for each plot. Predicted consumption was calculated by subtracting the predicted mass values of the Total 0-30cm post-burn linear model from the predicted mass values of the Total 0-30cm pre-burn linear model for each plot. We applied a linear model using the ‘lm’ R package to assess the relationship between observed and model predicted consumption values.

## Results

From the leaps and bounds regression, all linear models were significant (p<0.05) with an R^2^ ranging from of 0.65 to 0.74 using a subset of six variables and the exhaustive method of variable selection (Table 1, 2, Figure 3, 4). Using the pre-burn data, the best performing linear model was Total 0-30cm (R^2^ = 0.74, BIC = -29), while the worst performing model was FWD (R^2^ = 0.61, BIC = -12, Table 1). The pre- and post-burn data combined also illustrated a significant linear model, with an R^2^ of 0.69. The models including the post burn data, either alone or combined with pre-burn data, did not perform as well as the pre-burn data alone because of less variation in the mass values post-burn. Similarly, this was also true for FWD (Figure 4b). Furthermore, the linear relationship between observed and predicted consumption was significant (R^2^ = 0.70, *p* = 1.325 × 10^−11^, Fig.3d)

**Table 1.** Statistical results from leaps and bounds linear regression, which illustrate the resulting model selected subset of six TLS metrics used to predict the associated vegetation and fuel mass classes (linear models). All models had a df = 34, except Total 0-30cm Pre & Post was df = 75. See Appendix C for a full list of metrics and their definitions.

| **Vegetation & Fuel**<br>**Mass Class**<br>**(Linear Model)** | **Burn status** | **R<sup>2</sup>** | ***p*-value** | **RMSE** | **BIC** | **Model selected subset of six TLS metrics** |
| --- | --- | --- | --- | --- | --- | --- |
| Total | Pre | 0.72 | 4.5 x10<sup>-08</sup> | 228 | -24 | % PD2, Sk Prop Non-occluded2, Ku Prop Non-occluded2, % Space5, Mean Ht Voxels, Surface VD |
| Total no FWD | Pre | 0.65 | 1.3 x10<sup>-06</sup> | 208 | -17 | PD2, % PD2, PD5, Ht 65th Q, Ku Prop Non-occluded1, % Space3 |
| Total 0-30cm | Pre | 0.74 | 1. x10<sup>-08</sup> | 196 | -29 | % PD2, Sk Prop Non-occluded2, Ku Prop Non-occluded2, % Space5, Mean Ht Voxels, Surface VD |
| Total 0-30cm no FWD | Pre | 0.69 | 1.6 x10<sup>-07</sup> | 168 | -24 | Ht 90th Q, % PD 30<sup>th</sup> Q, Ku Prop Non-occluded1, SD Prop Non-occluded2, SD Ht Voxels, Ku CWD 1000 |
| FWD | Pre | 0.61 | 8.8 x10<sup>-06</sup> | 108 | -12 | SD Ht2, Sk Ht2, SD Ht5, 90th Q Ht, % Space5, Mean Ht Voxels |
| Fine Fuels | Pre | 0.71 | 5.9 x10<sup>-08</sup> | 148 | -25 | PD2, % PD2, Ht 30th Q, Ht 65th Q, % PD 10<sup>th</sup> Q, % PD 60<sup>th</sup> Q |
| Total 0-30cm Post | Post | 0.67 | 5.4 x10<sup>-07</sup> | 136 | -20 | Ht 45th Q, % PD 80<sup>th</sup> Q, % PD 90<sup>th</sup> Q, Max Tree Ht, Surface Mean Ht, Surface Sk Ht |
| Total 0-30cm Pre & Post | Pre & Post | 0.69 | 2.2 x10<sup>-16</sup> | 236 | -65 | Median Ht1, Ht 15th Q, % Occluded5, Mean Prop Non-occluded5, Mean Ht Voxels, Surface TGI |

**Table 2.**
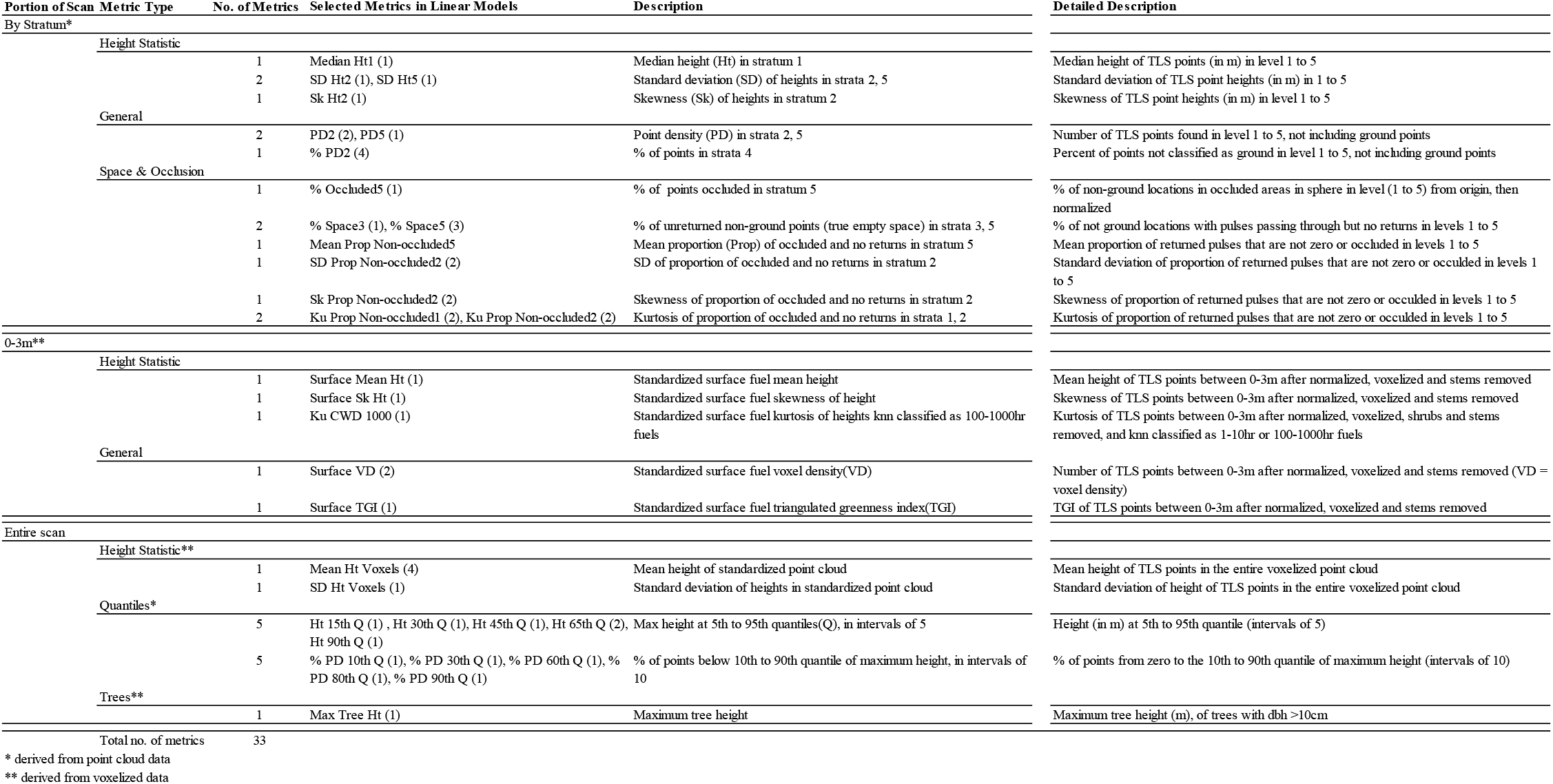
Selected metrics from linear models illustrating which portion of the scan in which the metric was derived, the metric type, as well as description of each metric. Stratum refers to the height layer within each scan in which the metric was derived, i.e., stratum 1: 0-0.5m, 2: 0.5-1m, 3: 1-1.5m, 4: 1.5-2m, and 5: >2m. The numbers in parentheses refer to the number of times that particular metric was selected across the eight linear models in this study. Metric type refers to either a height statistic, general or standard TLS metric (e.g. point density), metric by quantiles, or identified tree structures within each scan. A more detailed description of all metrics is found in Appendix C.

**Figure 3.**
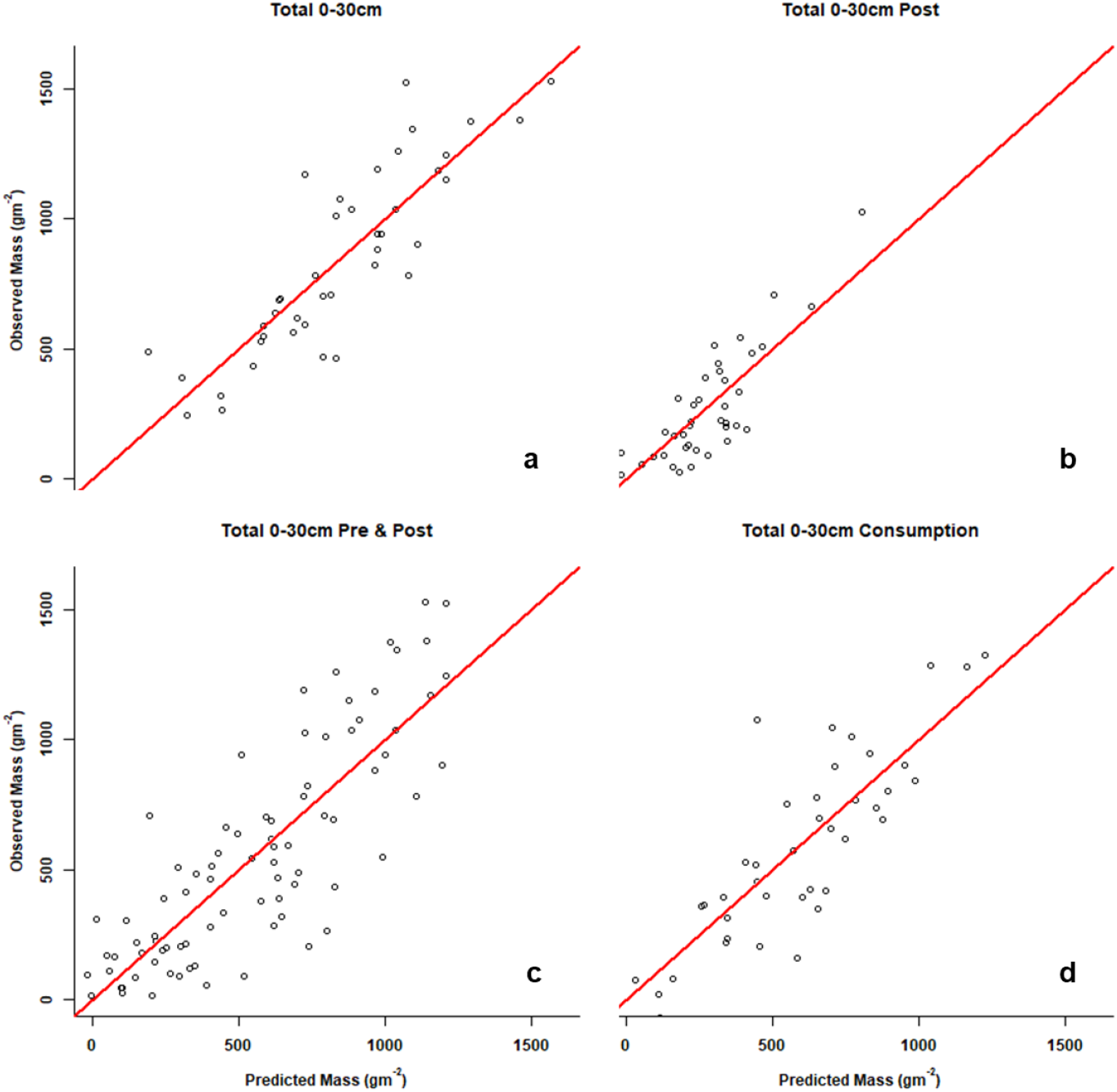
Observed vs. predicted surface biomass for the Total 0-30cm vegetation and fuel mass class, using the pre-burn (a, n=41), post-burn (b, n=41), pre- and post-burn combined c, (n=82), and consumption (d, n=41) linear models.

**Figure 4.**
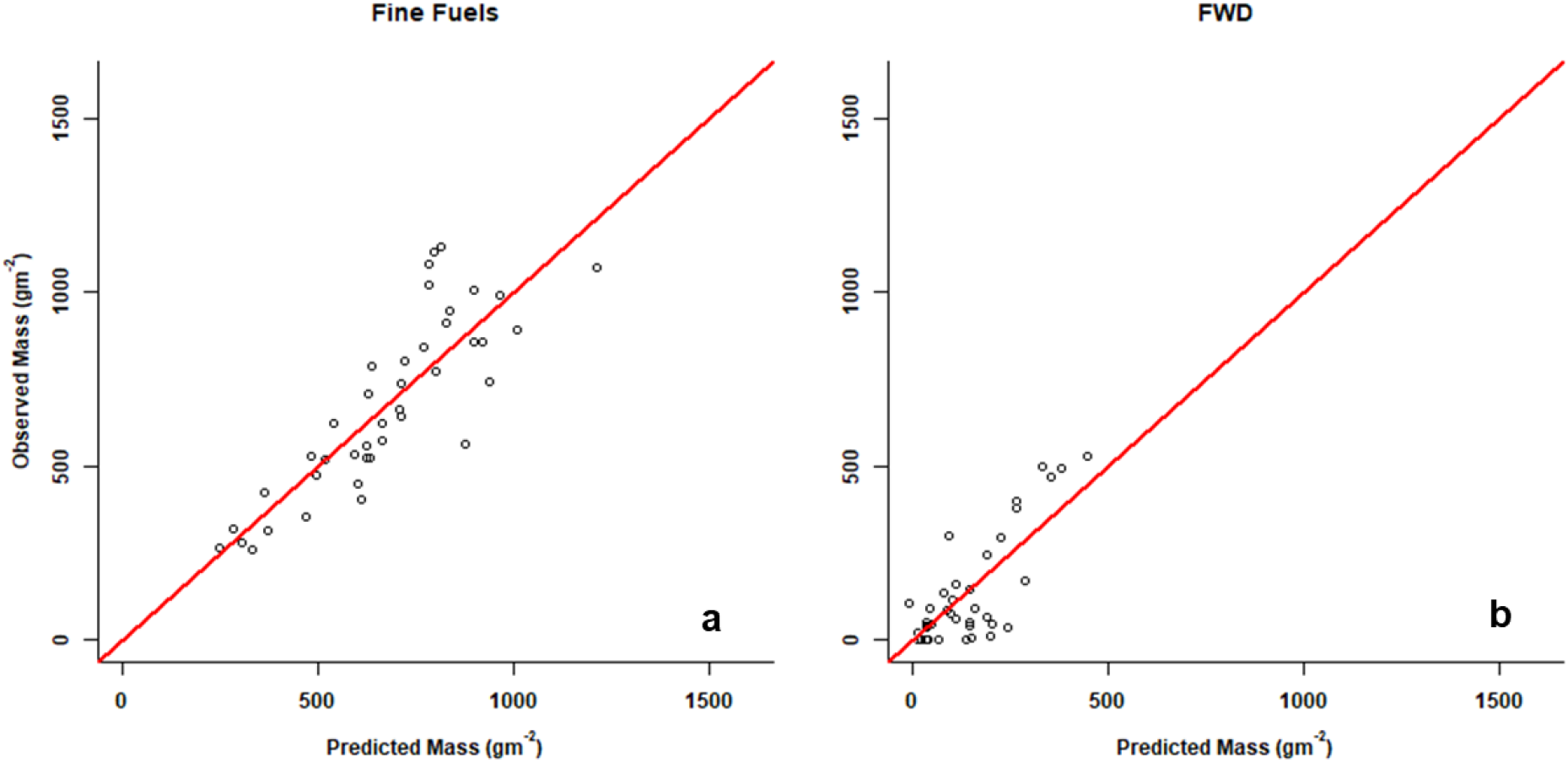
Observed vs. predicted surface biomass comparing the fine fuels (a) and fine woody debris (b) vegetation and fuel mass classes and their respective linear models.

These eight linear models with a possible six TLS metrics selected for each model resulted in 48 total selected metrics that were identified as most important in predicting mass (Table 1, 2). Descriptions of the selected metrics are in Table 2, and all metric descriptions are in Appendix C. The metrics were all significant predictors (*p* < 0.05). Though the metrics were not explicit between models; there were some similarities found. Thirty-three unique metrics, or 20% of the (162 total) input TLS metrics were selected among the eight models. Thirty (91% of 33) were from metrics calculated from different strata or height quantiles within each TLS scan. Of the remaining three, two included mean and standard deviation of heights in the entire scan. These two metrics were used in five of the eight models. The last one was a tree-metric: maximum tree height. This was only used once, in the Total 0-30cm Post model. Six metrics associated with space and occlusion (of 37 total) were selected within five models.

The selected metrics were identical for the Total and Total 0-30cm models (Table 1). Three metrics for the Total no FWD and Fine Fuels models were similar: PD2 and % PD2 representing the point density within 0.5-1m height profile and 65^th^ quantile in height within the scan. The selected metrics were unique between the Total 0-30cm and Total 0-30cm Post models. The metrics were more mixed in the Total 0-30cm Pre & Post model. This model had the largest range of metric types compared to the other models.

## Discussion

This study represents a critical step for operational application of single terrestrial laser scan methodology for measuring vegetation and fuel characteristics. We found that linear models can be used in frequently burned southeastern U.S. ecosystems to estimate fine-scale surface vegetation and fuel biomass using TLS vegetation metrics, and by predicting pre and post burn mass, they can estimate consumption. We also found that the first 95% of the total mass was found within the first 30 cm of the fuelbed and that using this value (e.g., Total 0-30cm) produced the highest performing linear models (e.g., Total 0-30cm, Table 1, Figure 3). This also includes estimating fine fuels, the largest proportion of mass pre-burn and mass consumed (Table 1, Figure 4a). FWD could also be estimated using this linear regression method (Figure 4b); however, we found that FWD represented 14% of the total mass; and, more importantly, was minimally reduced by the fire (i.e., no difference between pre and post-burn FWD) and thus did not contribute significantly to overall consumption. These findings illustrate a simple and practical method for predicting fine-scale mass from TLS-derived metrics.

The strength of the models that we developed can be attributed to our sample size, the large range of surface mass and fuel types sampled, and the variation in consumption during the fires. Furthermore, the three-dimensional nature of the TLS scans created a dataset from which a comprehensive list of TLS metrics could be derived and used to predict a selection of surface fuel characteristics that are known to drive fire behavior. Across the eight linear models, 33 of 162 TLS metrics (20% of total) were selected to best predict surface biomass. The selected metrics between models corresponded similarly where vegetation within each category was similar (e.g., Total vs. Total 0-30cm). The metrics were unique between the Total 0-30cm pre burn and Total 0-30cm post burn models, illustrating the distinct representation of surface vegetation structure before and after fire both within and across these scans. When combined (Total 0-30cm pre & post burn model), the metrics were dispersed between the individual models (Total 0-30cm vs. Total 0-30cm post), where complex similarities across the pre and post burn scans represented the higher range of variability between pre and post burn scans and associated biomass. Over 90% of the selected metrics (30 of 33) represented vegetation and space between vegetation among each scan’s height profile (Table 2, strata, quantiles, etc.), illustrating the profound complexity of vegetation at fine scales in these frequently burned systems. The three remaining metrics selected as important metrics were mean and standard deviation height of the entire voxelized point cloud as well max tree height. These three metrics were found in four models. Max tree height was only chosen in one model: the only post-scan model run (Total 0-30cm Post). This illustrates the significance of characterizing the entire forest structure across the vertical and horizontal plane (hence using height profiles and quantiles, etc., Table 2) beyond only explicit tree measurements to model the variability in surface vegetation and fuel biomass. Furthermore, empty space in the scan data were partitioned as gaps in vegetation and gaps due to occlusion. This was represented in 37 TLS of the total 162 metrics, of which six were chosen within five linear models (Table 1, 2). This illustrates the relevance of accounting for both true empty space and space that appears empty in scans but is an artifact of scan occlusion, all important factors when interpreting scan data and modeling surface biomass.

Given that surface fuel heterogeneity is complex and difficult to capture, the sensitivity of TLS is up to the task of capturing most of this fine-scale structural variation. Furthermore, our single-scan and linear modeling approach is relatively simple compared to other more direct one-to-one coupled methods. Other methods include linking fine-scale mass and volume of understory vegetation down to the 1.0 m^3^ and 6.25 × 10^−2^ m^3^ scale [27,32]. These methods use high-resolution TLS instrumentation that requires extensive processing to merge several TLS scans together and manually crop out each 3D vegetation plot or individual plant from the resulting merged 3D point cloud. Though this does create a more ‘complete’ 3D image of the vegetation by reducing occlusion and enhancing realism, its practical utility for extensive and repeated sampling work and management applications remains questionable. Furthermore, this method is likely prone to error associated with directly linking field measurements and TLS data cropped to the 1000 cm^3^ (or 10cm × 10cm × 10cm) scale because both are done manually and prone to error (e.g., wind during scans, placement of sampling frame, clipping biomass at 10cm intervals). Our field and processing approach was distinct from these methods by acquiring and processing one TLS scan from a push button instrument in one location (Pokswinski et al. 2021), then clipping *in situ*, drying, sorting, and weighing vegetation from only two strata (0-30, 30-100 cm), and running a linear regression technique that pulls out only a fraction (6 variables) of the TLS metrics to explain up to 74% of the variation in surface vegetation and fuel biomass. Even if we only measured total dry mass, without sorting into explicit vegetation and fuel categories, or even height categories (including FWD), our linear models still performed well (e.g., ‘Total’ category, R^2^=0.70, Table 1) and could be readily tested in other systems.

The TLS instrument used in this study was useful because of its simple and non-bias push-button approach. It could, however, prove useful to test the relationship between metrics derived from other TLS instruments and surface biomass to expand this study’s applicability. For instance, there are other 360° scanners that have different frequencies and ranges [39]. There are mobile terrestrial scans that, while walking a transect for example, could collect and process a more complete 3D point cloud of vegetation structure by simultaneously eliminating the merging process required for stationary TLS scans and reducing issues of occlusion. Though compared to our single-scan approach, these mobile scanners are currently more expensive, more difficult to operate, and quality of the point-cloud is dependent on the pace and path taken by the user, which limits replicability [34]. This single-scan approach can be repeated in the same plot, as we did for consumption, for long-term consistent data collection with minimal bias.

Creating TLS-derived models that estimate variation in surface biomass within forested stands pave the way for creating realistic inputs and robust test datasets for spatially explicit fire behavior models (Parsons et al 2018). These data are particularly useful in ecosystems dominated by fine scale variation in surface fuels (Hiers et al. 2009) and for inputs into high resolution coupled fire-atmosphere models (Linn et al. 2001, Mell et al. 2009, Linn et al. 2020) that operate in three dimensions with the ability to represent within-stand variability (Hiers et al. 2020). Such high-resolution representation of fuel variation will improve our prediction of heterogeneity in fire effects and the underlying physical influences of vegetation on the fire environment [13]. In this study, we found that the variability in surface mass across these two burn units (658 ha total) was considerable (mean: 841 g m^-2^, stdev: 350 g m^-2^ from the ‘Total 0-30’ model (Figure 2), highlighting the need to think beyond stand level averages [66]. Consumption followed this same trend (mean: 580 g m^-2^, std: 353 g m^-2^), allowing for connections of energy-released-by-fire to fire effects (O’Brien et al. 2018). Both results demonstrate the ability to represent 3D variability in mass and consumption across a stand as well as the significance of low (here <30cm) vegetation, particularly fine-fuels, when coarse wood consumption is negligible. As such, we recommend these TLS data for pre-burn biomass estimates as inputs to coupled fire-atmosphere models, while our consumption estimates can be compared to model outputs.

This single scan sampling technique was designed to be simple and efficient, specifically for long-term monitoring [44]. Each scan captures a high-resolution 3D image of a plot that can be analyzed repeatedly as new analyses and metrics are developed with minimal to no post-processing. Incorporating surface biomass estimation into this approach is critical particularly for frequently burned ecosystems where mass varies at fine-scales and influences fire behavior and effects at these same scales [15]. Broader stand-level averages do not account for this fine-scale heterogeneity and are not sufficient for coupling to fire-atmosphere models and projections of resulting fire effects [10,67]. This approach can add more explicit estimates of mass to existing broad-scale approaches, such as ecosystem-level photo-load series and custom fuel models [12,68]. The benefit for management, especially if preliminary dry mass values have been incorporated, is that scans can be taken efficiently within vegetation types of interest and run through the established linear models to provide explicit variability of surface biomass found within and across stands. Surface mass can be quite variable across stands, even within the same ecosystem type (e.g., longleaf pine Flatwoods vs sandhills) based on fire histories, time-since-fire, management practices, and general site characteristics (soil, overstory density, etc.). This approach could be used for fire effects monitoring associated with surface biomass change detection, both before and after a fire, monitoring long-term restoration and maintenance of target surface fuel characteristics over time.

Future work for this study includes relevance across similar sites and other frequently burned ecosystems. The extension of these models will require testing across other regional longleaf pine forests or frequently burned ecosystems in other regions. We expect similarities in ecosystems with comparable physiognomy, though model-selected TLS metrics may vary depending on local factors, such as dominant trees, understory and site characteristics (e.g. soil, elevation, topography, climate, management). Validation procedures are needed to confirm their utility in prediction of fine-scale variability in surface mass used for fire behavior research, particularly using these fuels predictions, coupled with site characteristics to create 3D surface fuel maps. Coupling these TLS derived surface fuel biomass can be combined with airborne laser scanning to provide full spatially explicit vegetation distribution across a given site.

In conclusion, this study illustrates an approach to expand the utility of single location terrestrial laser scanning for prescribed fire management and monitoring fire effects associated with surface vegetation and fuel biomass. This study demonstrates the capability of simple TLS methods to measure within and across stand variability as well as estimated consumption. Lastly, these data and the linear models presented here could prove useful for deriving inputs to and validating outputs from coupled fire-atmosphere models, particularly for prescribed fire research and management.

## Acknowledgements

This project was funded in part by the Department of Defense’s (DoD) Strategic Environmental and Research Development Program projects RC19-1119 and RC20-1346 and supported by DoD’s Environmental Security Technology Certification Program project: the Integrated Research Management Team, with special thanks to James Furman for his profound leadership. We give an extra special thanks to the Ft. Stewart Natural Resources and Forestry Staff led by Bryan Whitmore. They illustrated exemplary support during this field campaign, particularly in facilitating the safety of our field crew, field support, and executing the experimental burns with extreme professionalism and efficiency. We thank the Southern and Northern Research Stations for their support, particularly the Athens Prescribed Fire Laboratory of the Center for Forest Health and Disturbance in Athens, GA. We thank Tall Timbers Research Station for their support of technical staff. We thank field preparation, sampling and laboratory efforts provided by Derek Wallace, C. Wade Ross, Greg Chapman, Vanessa Niemczyk, Jacob Ney, Irenee Payne, and Quinn Hiers.

