## Appendices for "Terrestrial laser scan metrics predict surface vegetation biomass and consumption in a frequently burned southeastern U.S. ecosystem"

**Appendix A.** Table of original vegetation or fuel categories collected in the field and their descriptions.

| Vegetation or Fuel Category | Description |
|---|---|
| Woody Live | Live material from evergreen and deciduous broadleaf shrubs or trees aboveground (i.e., stems, leaves, flowers, buds, etc.) |
| Now Dead Woody Vegetation | Only in post-burn sampling to classify pre-burn woody live stems that were partially consumed by the prescribed fire and the aboveground plant was clearly dead (aka top-killed) |
| Woody Litter | Downed leaf and litter material from evergreen and deciduous broadleaf shrubs or trees detached from its source (i.e., leaves, flowers, buds, etc.) |
| 1-hr | Downed dead branches, twigs, and other small woody pieces that are severed from their original source of growth, and dead woody species that is still standing and attached to the ground and is less than 0.25 inch (1.27 cm) in diameter |
| 10-hr | Downed, dead branches, twigs, and other small woody pieces that are severed from their original source of growth, female cones (i.e., megastrobilus, seed cone, or ovulate cone) from non-*Pinus* species, and dead woody species that are still standing and attached to the ground and is 0.25 inch to 1.0 inch (1.27 to 2.54 cm) in diameter |

| | |
|---|---|
| 100-hr | Downed, dead tree and shrub boles, large limbs, and other woody pieces that are severed from their original source of growth and dead woody species that is still standing and attached to the ground and is 1.0 inch to 3.0 inch (2.54 to 7.6 cm) in diameter |
| 1000-hr | Downed, dead tree and shrub boles, large limbs, and other woody pieces that are severed from their original source of growth and is 3.0 inch to 8 inch (7.6 cm to 20.3 cm) in diameter. Note that no 1000-hr fuels were found in our plots for this study. |
| Pinecones | Intact female cones (i.e., megastrobilus, seed cone, or ovulate cone) from *Pinus* species |
| Conifer Litter | Needle from conifers other than Pinus species and downed woody material from conifer species that is too small to fit into the 1-hour fuel category (ex: paper-thin pieces of bark, male pollen cones (aka microstrobilus), and pinecone fragments) |
| Pine Needles | Downed needles from *Pinus* species with long or short needles |
| Fine Vegetation | Live and dead material from bunchgrass species, wiregrass species, other graminoids, forbs, vines, and conifer seedlings |

**Appendix B.** Surface biomass values (g $m^{-2}$) for each plot (row, n=41) corresponding to the vegetation and fuel mass classes (headings) used in this study and described in the text. All values represent biomass found before the fire (pre-burn), except for the last column (Total 0-30cm Post) that is the biomass measured after the fire in the post-burn plots. 'Total 0-30cm' and 'Total 0-30cm Post' classes were combined (n=82) to produce the 'Total 0-30cm Pre & Post' category (not shown here).

| **Total** | **Total no FWD** | **Total 0-30cm** | **Total 0-30cm no FWD** | **Fine Fuels** | **FWD** | **Total 0-30cm Post** |
|---|---|---|---|---|---|---|
| 318.6 | 278.2 | 318.6 | 278.2 | 278.2 | 40.4 | 84.08 |
| 1124.92 | 1011.4 | 1075.04 | 961.52 | 945.96 | 113.52 | 178.92 |
| 470.16 | 432.36 | 466.08 | 428.28 | 404.44 | 37.8 | 43.52 |
| 942 | 854.8 | 942 | 854.8 | 854.8 | 87.2 | 101.12 |
| 787.68 | 787.68 | 782.2 | 782.2 | 787.68 | 0 | 16.76 |
| 490.68 | 472.96 | 490.68 | 472.96 | 472.96 | 17.72 | 95.16 |
| 553.2 | 531.76 | 550.32 | 528.88 | 521.68 | 21.44 | 23.84 |
| 565.28 | 320.72 | 565.28 | 320.72 | 320.72 | 244.56 | 110.4 |
| 622.16 | 533.24 | 618.72 | 529.8 | 521.96 | 88.92 | 219.12 |
| 739.16 | 725.88 | 685.72 | 672.44 | 555.36 | 13.28 | 169.84 |
| 1396.52 | 1020.12 | 1341.52 | 965.12 | 911.12 | 376.4 | 57.64 |
| 529.08 | 529.08 | 529.08 | 529.08 | 527.24 | 0 | 507.92 |
| 853.08 | 848.04 | 783.64 | 778.6 | 739.48 | 5.04 | 46.16 |
| 1122.36 | 630.76 | 1007.96 | 516.36 | 449.56 | 491.6 | 310.08 |
| 1373.76 | 876.28 | 1373.76 | 876.28 | 856.16 | 497.48 | 92.8 |
| 1318.32 | 1281.68 | 1150.44 | 1113.8 | 1003.8 | 36.64 | 207.16 |
| 1727.36 | 1434.68 | 1526.12 | 1233.44 | 1077.72 | 292.68 | 201.56 |
| 1648 | 1600.6 | 1256.72 | 1209.32 | 1020.44 | 47.4 | 213.04 |
| 467.52 | 406.36 | 461.44 | 400.28 | 353.04 | 61.16 | 302.6 |
| 1178.92 | 1131.12 | 1167.08 | 1119.28 | 1131.12 | 47.8 | 91.6 |
| 1276.64 | 806.28 | 1188.64 | 718.28 | 621.92 | 470.36 | 414.28 |
| 756.2 | 620.32 | 702.24 | 566.36 | 620.32 | 135.88 | 129.8 |
| 1512.44 | 983.6 | 1378.16 | 849.32 | 772.16 | 528.84 | 1027.36 |
| 434.6 | 434.6 | 434.6 | 434.6 | 423.52 | 0 | 122.16 |
| 591.92 | 527.72 | 591.92 | 527.72 | 516.24 | 64.2 | 386.96 |
| 266.16 | 266.16 | 243.44 | 243.44 | 266.16 | 0 | 165.04 |
| 913.56 | 612.6 | 900 | 599.04 | 564.56 | 300.96 | 483.2 |
| 694.28 | 642.32 | 693.24 | 641.28 | 642.32 | 51.96 | 332.16 |
| 881.68 | 737.24 | 881.68 | 737.24 | 737.24 | 144.44 | 661.72 |
| 495.8 | 495.8 | 389.88 | 389.88 | 314.28 | 0 | 146 |
| 614.36 | 572.28 | 586.64 | 544.56 | 572.28 | 42.08 | 224.88 |
| 820.84 | 660.92 | 820.84 | 660.92 | 660.92 | 159.92 | 205.64 |
| 707.52 | 707.52 | 707.52 | 707.52 | 707.52 | 0 | 15.4 |

| | | | | | | |
|---|---|---|---|---|---|---|
| 1575.24 | 1175.08 | 1520.8 | 1120.64 | 1114.44 | 400.16 | 512.4 |
| 1217.8 | 1197.28 | 1033.44 | 1012.92 | 800.76 | 20.52 | 279.44 |
| 1244.08 | 1074.24 | 1244.08 | 1074.24 | 1068.6 | 169.84 | 442.52 |
| 1212.76 | 1123 | 1185.44 | 1095.68 | 988.28 | 89.76 | 282.24 |
| 1134.36 | 1060.12 | 1037.4 | 963.16 | 842.68 | 74.24 | 378.04 |
| 938.92 | 889.4 | 938.92 | 889.4 | 889.4 | 49.52 | 544.56 |
| 639.04 | 531.76 | 639.04 | 531.76 | 531.76 | 107.28 | 709.2 |
| 264.04 | 264.04 | 264.04 | 264.04 | 261.72 | 0 | 190.8 |

**Appendix C.** The 162 TLS metrics used in the linear models for this study. The gray rows are those chosen in our regression models for this study; see Table 1. These illustrate which portion of the scan the metric was derived, the metric type, as well as a description of each metric. Stratum refers to the height layer within each scan in which the metric was derived, i.e., strata 1: 0-0.5m, 2: 0.5-1m, 3: 1-1.5m, 4: 1.5-2m, and 5: >2m. 'Metric type' refers to either a height statistic, general or standard TLS metric (e.g. point density), metric associated with space and occlusion of TLS points within the scan (e.g. true empty space, occlusion by trees), metric by height quantiles, or identified tree structure within each scan. Metrics were calculated from either the point cloud or voxelization of the point cloud to 2cm x 2cm x 2cm voxels. "Standardized" means the point cloud was voxelized before the metric was calculated. Standardized surface fuels means that identified tree stems were removed before voxelization of each point cloud below 3m in height. All scans were normalized to account for topographic variation and cropped to a 15m radius from center before any metric calculation. Triangular Greenness Index (TGI) = ((Green -0.39)*(Red - 0.61))*Blue values extracted from the BLK camera. Visible Atmospherically Resistant Index (VWRI) = (Green-Red)(Green+Red-Blue) extracted from the BLK camera. See text for more details of the methodology.

| Portion of Scan | Metric type | Voxelized/ Point Cloud | No. of Metrics | Metric | Description |
|---|---|---|---|---|---|
| By stratum | General | Point Cloud | 5 | PD(1 to 5), e.g., PD1 | Point density (PD) in strata 1 to 5 |
| By stratum | General | Point Cloud | 5 | % PD(1 to 5) | % of points in strata 1 to 5 |
| By stratum | General | Point Cloud | 5 | TGI(1 to 5) | Triangular Greenness Index (TGI) in strata 1 to 5 |
| By stratum | General | Point Cloud | 5 | VARI(1 to 5) | Visual Atmospheric Resistance Index (VARI) in strata 1 to 5 |
| By stratum | Height statistic | Point Cloud | 5 | Mean Ht(1 to 5) | Mean height (Ht) in strata 1 to 5 |
| By stratum | Height statistic | Point Cloud | 5 | Median Ht(1 to 5) | Median height in strata 1 to 5 |
| By stratum | Height statistic | Point Cloud | 5 | SD Ht(1 to 5) | Standard deviation (SD) of heights in strata 1 to 5 |
| By stratum | Height statistic | Point Cloud | 5 | Sk Ht(1 to 5) | Skewness of heights (Sk) in strata 1 to 5 |
| By stratum | Height statistic | Point Cloud | 5 | Ku Ht(1 to 5) | Kurtosis (Ku) of heights in strata 1 to 5 |
| By stratum | Space & Occlusion | Point Cloud | 5 | % Occluded(1 to 5) | % of non-ground points occluded in strata 1 to 5 |

| | | | | | |
|---|---|---|---|---|---|
| By stratum | Space & Occlusion | Point Cloud | 5 | % Space(1 to 5) | % of unreturned non-ground points (true empty space) in strata 1 to 5 |
| By stratum | Space & Occlusion | Point Cloud | 5 | Mean Prop Non-occluded(1 to 5) | Mean proportion(Prop) of occluded and no returns in strata 1 to 5 |
| By stratum | Space & Occlusion | Point Cloud | 5 | SD Prop Non-occluded(1 to 5) | SD of proportion of occluded and no returns in strata 1 to 5 |
| By stratum | Space & Occlusion | Point Cloud | 5 | Sk Prop Non-occluded(1 to 5) | Sk of proportion of occluded and no returns in strata 1 to 5 |
| By stratum | Space & Occlusion | Point Cloud | 5 | Ku Prop Non-occluded(1 to 5) | Ku of proportion of occluded and no returns in strata 1 to 5 |
| 0-3m | General | Voxelized | 2 | PD CWD (10 or 1000) | Standardized surface fuel PD classified as 1-10hr or 100-1000hr fuels |
| 0-3m | General | Voxelized | 2 | TDI CWD (10 or 1000) | Standardized surface fuel TDI classified as 1-10hr or 100-1000hr fuels |
| 0-3m | General | Voxelized | 2 | VARI CWD (10 or 1000) | Standardized surface fuel VARI classified as 1-10hr or 100-1000hr fuels |
| 0-3m | Height statistic | Voxelized | 2 | Mean CWD (10 or 1000) | Standardized surface fuel mean height classified as 1-10hr or 100-1000hr fuels |
| 0-3m | Height statistic | Voxelized | 2 | Median CWD (10 or 1000) | Standardized surface fuel median height classified as 1-10hr or 100-1000hr fuels |
| 0-3m | Height statistic | Voxelized | 2 | SD CWD (10 or 1000) | Standardized surface fuel SD of height classified as 1-10hr or 100-1000hr fuels |
| 0-3m | Height statistic | Voxelized | 2 | Sk CWD (10 or 1000) | Standardized surface fuel Sk of height classified as 1-10hr or 100-1000hr fuels |
| 0-3m | Height statistic | Voxelized | 2 | Ku CWD (10 or 1000) | Standardized surface fuel Ku of height classified as 1-10hr or 100-1000hr fuels |
| Entire scan | General | Point Cloud | 1 | Ground PD | Number of TLS points classified as ground |
| Entire scan | General | Point Cloud | 1 | Veg PD | Number of TLS points not classified as ground |
| Entire scan | General | Point Cloud | 1 | % Ground | % of points classified as ground |
| Entire scan | General | Point Cloud | 1 | TGI | TGI |
| Entire scan | General | Point Cloud | 1 | VARI | VARI |
| Entire scan | General | Point Cloud | 1 | % Above Mean Ht | % of non-ground points above mean height |
| Entire scan | General | Point Cloud | 1 | % Above 2SD Mean Ht | % of non-ground points 2SD above mean height |
| Entire scan | General | Voxelized | 1 | Total Volume | PD of standardized point cloud |
| Entire scan | General | Voxelized | 1 | TGI Voxels | TGI of standardized point cloud |

| | | | | | |
|---|---|---|---|---|---|
| Entire scan | General | Voxelized | 1 | VARI Voxels | VARI of standardized point cloud |
| Entire scan | Height statistic | Point Cloud | 1 | Maximum Ht | Maximum height of TLS points in the entire scan |
| Entire scan | Height statistic | Point Cloud | 1 | Mean Ht | Mean height of TLS points in the entire scan |
| Entire scan | Height statistic | Point Cloud | 1 | SD Ht | SD of TLS point heights in the entire scan |
| Entire scan | Height statistic | Point Cloud | 1 | Sk Ht | Sk of TLS point heights in the entire scan |
| Entire scan | Height statistic | Point Cloud | 1 | Ku Ht | Ku of TLS point heights in the entire scan |
| Entire scan | Height statistic | Voxelized | 1 | Mean Ht Voxels | Mean height of standardized point cloud |
| Entire scan | Height statistic | Voxelized | 1 | Median Ht Voxels | Median height of standardized point cloud |
| Entire scan | Height statistic | Voxelized | 1 | SD Ht Voxels | SD of heights in standardized point cloud |
| Entire scan | Height statistic | Voxelized | 1 | Sk Ht Voxels | Sk of heights in standardized point cloud |
| Entire scan | Height statistic | Voxelized | 1 | Ku Ht Voxels | Ku of heights in standardized point cloud |
| Entire scan | Space & Occlusion | Point Cloud | 1 | % Total Unreturned Points | % of unreturned points overall |
| Entire scan | Space & Occlusion | Point Cloud | 1 | % Area Occluded Points | % of possible non-ground points from occluded |
| Entire scan | Space & Occlusion | Point Cloud | 1 | % True Open Space | % of unreturned non-ground points (true empty space) |
| Entire scan | Space & Occlusion | Point Cloud | 1 | Mean Prop Non-occluded | Mean proportion of true points that are not occluded |
| Entire scan | Space & Occlusion | Point Cloud | 1 | SD Prop Non-occluded | SD of proportion of occluded and no returns in entire scan |
| Entire scan | Space & Occlusion | Point Cloud | 1 | Sk Prop Non-occluded | Sk of proportion of occluded and no returns in entire scan |
| Entire scan | Space & Occlusion | Point Cloud | 1 | Ku Prop Non-occluded | Ku of proportion of occluded and no returns in entire scan |
| Entire scan | Quantiles | Point Cloud | 19 | Ht 5th Q to Ht 95th Q | Height at 5th to 95th quantiles in intervals of 5 |

| | | | | | |
|---|---|---|---|---|---|
| Entire scan | Quantiles | Point Cloud | 9 | % PD 10th Q to % PD 90th Q | % of points below 10th to 90th quantile of max ht in intervals of 10 |
| Entire scan | Trees | Voxelized | 1 | Total BA | Total basal area |
| Entire scan | Trees | Voxelized | 1 | Mean BA | Mean basal area |
| Entire scan | Trees | Voxelized | 1 | Mean Tree Ht | Mean tree height |
| Entire scan | Trees | Voxelized | 1 | Mean DBH | Mean diameter at breast height (DBH) of all detected trees >4cm DBH |
| Entire scan | Trees | Voxelized | 1 | No. Trees | Number of trees detected |
| Entire scan | Trees | Voxelized | 1 | Max Tree Ht | Maximum tree height |
| Entire scan | Trees | Voxelized | 1 | SD Tree Hts | SD of tree heights |
| Entire scan | Trees | Voxelized | 1 | Mean Canopy Base Ht | Mean tree canopy base height |
| 0-3m | General | Voxelized | 1 | Surface VD | Standardized surface fuel VD |
| 0-3m | General | Voxelized | 1 | Surface TGI | Standardized surface fuel TGI |
| 0-3m | General | Voxelized | 1 | Surface VARI | Standardized surface fueVARI |
| 0-3m | Height statistic | Voxelized | 1 | Surface Mean Ht | Standardized surface fuel mean height |
| 0-3m | Height statistic | Voxelized | 1 | Surface Median Ht | Standardized surface fuel median height |
| 0-3m | Height statistic | Voxelized | 1 | Surface SD ht | Standardized surface fuel SD of height |
| 0-3m | Height statistic | Voxelized | 1 | Surface Sk ht | Standardized surface fuel Sk of height |
| 0-3m | Height statistic | Voxelized | 1 | Surface Ku ht | Standardized surface fuel Ku of height |
